# An mTOR-Tfeb-Fabp7a signaling axis can be harnessed to ameliorate *bag3* cardiomyopathy in adult zebrafish

**DOI:** 10.1101/2024.10.24.620101

**Authors:** Yonghe Ding, Feixiang Yan, Baul Yoon, Wei Wei, David Mondaca Ruff, Yuji Zhang, Xueying Lin, Xiaolei Xu

## Abstract

Dysregulated proteostasis in cardiomyocytes is an important pathological event in *BAG3* cardiomyopathy, which can be repaired by inhibiting mechanistic target of rapamycin (mTOR) for cardioprotective effects. Here, we aimed to uncover additional pathological events and therapeutic target genes via leveraging zebrafish genetics. We first assessed transcription factor EB (*tfeb*), a candidate gene that encodes a direct downstream phosphorylation target of mTOR signaling. We found that cardiomyocyte-specific transgenic overexpression of *tfeb* (*Tg[cmlc2:tfeb]*) is sufficient to repair defective proteostasis, attenuate accelerated cardiac senescence, a previously unrecognized phenotype in the *bag3* cardiomyopathy model, and rescue cardiac dysfunction. Next, we compared cardiac transcriptomes between the *Tg(cmlc2:tfeb)* transgenic fish and the *mtor^xu015/+^* mutant, and tested 4 commonly downregulated lipodystrophy genes using an F0-based genetic assay. We found that inhibition of the fatty acid binding protein a (*fabp7a*) gene, but not the other 3 genes, exerts therapeutic effects on *bag3* cardiomyopathy. Conversely, *fabp7a* expression is elevated in *bag3* cardiomyopathy model and cardiomyocyte-specific overexpression of *fabp7a* resulted in dysregulated proteostasis, accelerated cardiac senescence, as well as cardiac dysfunction. Together, these genetic studies in zebrafish uncovered Fabp7a activation and accelerated cardiac senescence as important pathological events in *bag3* cardiomyopathy. The mTOR-Tfeb-Fabp7a signaling axis can be harnessed to repair these pathological changes and exert cardioprotective effects.

## Introduction

Dysregulated proteostasis in cardiomyocytes is an important pathological event in cardiomyopathies that further drives the pathogenesis ^1^. This notion is supported by the discovery of *BAG3* as a causative gene for dilated cardiomyopathies (DCM) ^2–4^. *BAG3* encodes a co-chaperone protein that binds to heat shock protein 70 and plays an important role in proteostasis, especially in the turnover of sarcomeric proteins ^5^. Our recent study of a *bag3^e^*^2^*^/e^*^2^ cardiomyopathy model in zebrafish further suggested that repairing dysregulated proteostasis can be an effective therapeutic strategy: inhibition of the mechanistic target of rapamycin (mTOR) through an *mtor* haploinsufficiency mutant (*mtor^xu01^*^5^*^/+^*) exerted cardioprotective effects ^6^. However, the downstream gene targets and signaling pathways by which inhibition of mTOR signaling confers cardioprotective effects remain to be elucidated. As a direct downstream phosphorylation target of mTOR signaling (through phosphorylation at S211 in the human TFEB protein), mammalian TFEB (Transcription factor EB) is a master transcriptional regulator of lysosomal biogenesis and autophagy genes and has been implicated in a diverse of functions such as nutrient sensing, lipid catabolism and protein homeostasis ^7, 8^. Overexpression of *TFEB* has been found to exert therapeutic benefits to neurodegenerative diseases ^9^ and to repair CryABR^120G^ overexpression-induced cardiac proteinopathy ^10^. Thus, *tfeb* is a promising downstream candidate gene that could recapitulate the cardioprotective effects of mTOR inhibition on the *bag3^e^*^2^*^/e^*^2^ cardiomyopathy model.

Like dysregulated proteostasis, accelerated cardiac senescence has also been reported to contribute to the pathogenesis of some cardiomyopathy models such as anthracycline-induced cardiotoxicity (AIC) ^11^. Both dysregulated proteostasis and increased cellular senescence have been found as one of the twelve hallmarks of normative aging ^12^. While the expression of Bag3 protein has been linked to aging, as indicated by a significantly elevated ratio of Bag3/Bag1 in neurons of aged rodent brain ^13^, whether cellular senescence contributes to the pathogenesis of *bag3^e^*^2^*^/e^*^2^ cardiomyopathy remains unestablished.

Fatty acid binding proteins (FABP) are a family of intracellular lipid-binding proteins mostly known for serving as lipid chaperones that mediate lipid transportation and lipid metabolic homeostasis. In humans, at least 10 FABPs have been identified and categorized according to their expression levels in various tissues and organs ^14^. Several FABP family members, such as FABP3, FABP4, and FABP9, have been identified as potential downstream target genes of TFEB, playing roles in cellular lipid metabolic process ^15^. FABP3, also known as the heart-specific FABP, is most abundantly expressed in the heart and cardiac muscle, emerging as a promising biomarker for coronary and peripheral artery disease ^16^. In contrast, FABP7 is known as the brain-type FABP, which mainly functions in brain development, learning and memory, sleeping disorder, and tumorigenesis ^17, 18^. There are no reports on either expression or functions of Fabp7 in the heart or cardiac diseases.

Here, we aim to leverage the efficient genetic tools offered by zebrafish to decipher the pathogenesis of *bag3^e^*^2^*^/e^*^2^ cardiomyopathy and the signaling axis that confers the cardioprotective effects of *mtor* inhibition. We found that cardiomyocyte-specific overexpression of *tfeb* is sufficient to recapitulate the cardioprotective effects of *mtor* inhibition. We then searched for additional modifying genes by testing differentially expressed genes suggested by a transcriptome comparison between the *mtor* inhibition and the *tfeb* transgenic overexpression fish lines. Using an F0-based genetic assay, we identified previously unreported expression of Fabp7a in the heart that is aberrantly activated in the *bag3* cardiomyopathy model. Lastly, Fabp7a inhibition recapitulates the cardioprotective effects of *mtor* inhibition and *tfeb* overexpression.

## Results

### 1. Cardiomyocyte-specific overexpression of *tfeb* recapitulated the cardioprotective effects of *mtor^xu0^*^15^*^/+^* in the *bag3* cardiomyopathy model

We recently reported that a frameshift mutation in the 2nd exon of the zebrafish *bag3* gene resulted in cardiomyopathy-like phenotypes at 6 months of age, termed *bag3^e^*^2^*^/e^*^2^ cardiomyopathy ^6^. A zebrafish mechanistic target of rapamycin (mTOR) haploinsufficiency mutant (*mtor^xu0^*^15^*^/+^*) exerted cardioprotective effects. Because Tfeb is a direct downstream phosphorylation target of mTOR signaling, we investigated whether the cardioprotective effects of mTOR inhibition can be recapitulated by Tfeb overexpression. We tested *Tg(cmlc2:tfeb),* a transgenic fish line with cardiomyocyte-specific overexpression of *tfeb* ^19^, and found that the *Tg(cmlc2:tfeb)* transgene effectively rescued cardiomyopathy-like phenotypes including increased ventricular chamber size, damaged myocardium structure, reduced cardiac function, shortened lifespan, and re-activation of fetal gene programs in the *bag3^e^*^2^*^/e^*^2^ mutant fish **(Fig. 1A-F**). Next, we performed a Western blot experiment and found that dysregulated autophagy and proteostasis in the *bag3^e^*^2^*^/e^*^2^ cardiomyopathy hearts, as indicated by an aberrant response of LC3 II to bafilomycin A1 (BafA1) and elevated ubiquitinated protein aggregations, were partially restored by the *Tg(cmlc2:tfeb)* transgene (**Fig. 1G and H**). Collectively, these data suggest that cardiomyocyte-specific overexpression of *tfeb* recapitulated the therapeutic effects of mTOR inhibition in the *bag3* cardiomyopathy model.

**Fig. 1.**
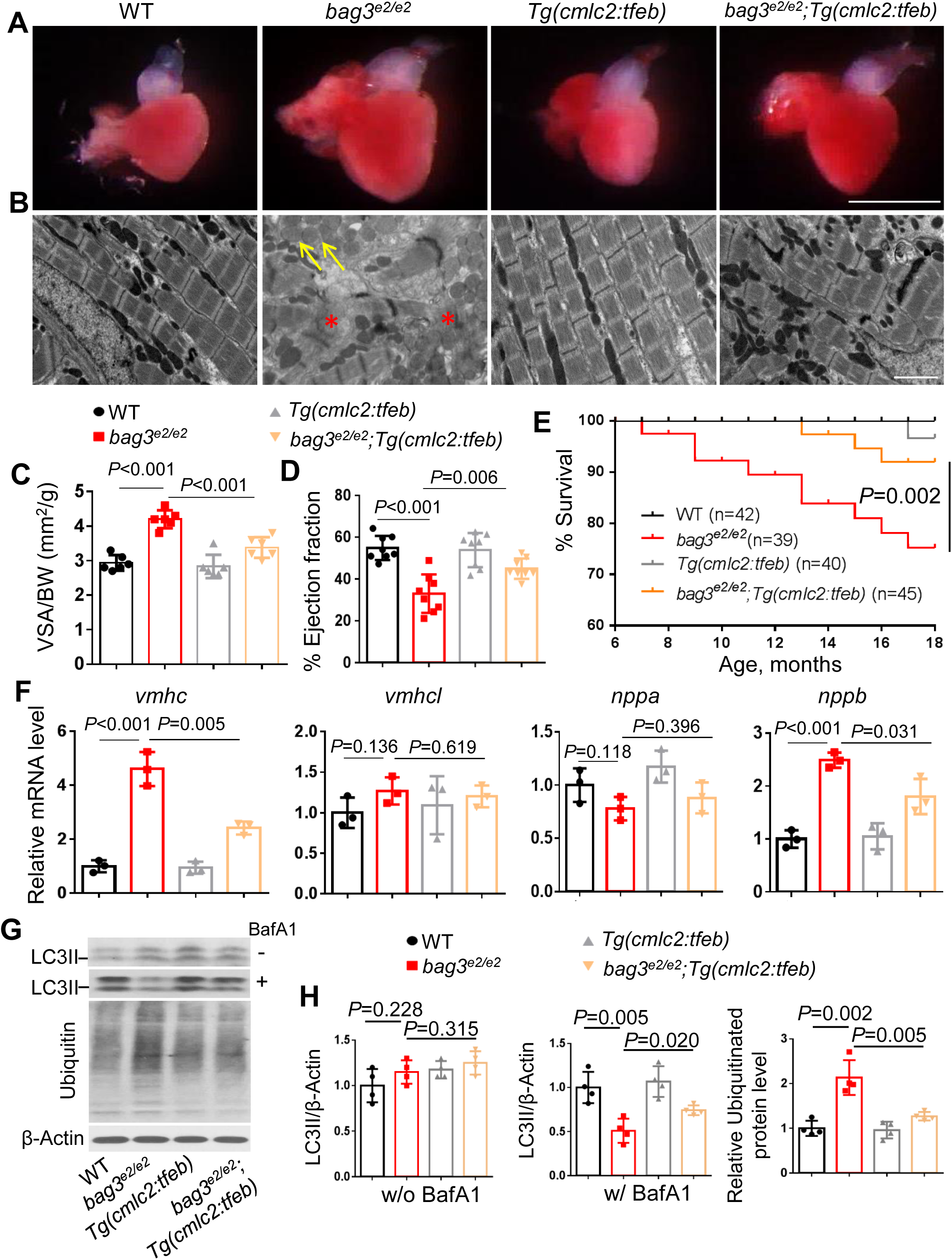
Cardiomyocyte-specific overexpression of *tfeb* alleviated *bag3* cardiomyopathy phenotypes and restored impaired proteostasis. **A-C,** Bright-field images of dissected hearts (A), confirmative images of transmission electron microscope (TEM) (B) and quantification analysis (C) show enlarged ventricular surface area (VSA) normalized to body weight (BW), impaired sarcomere structure (Asterisks), and mitochondrial swelling (Arrows) phenotypes in the *bag3^e2/e2^* mutant hearts, which were ameliorated by *Tg(cmlc2:tfeb),* a cardiomyocyte-specific *tfeb* overexpression transgenic line. Scale bars in A, 1 mm; in B, 20 μm. **D-E,** Ejection fraction (EF) (D) and Survival (in %) (E) of *bag3^e2/e2^;Tg(cmlc2:tfeb)* double mutant/transgenic fish compared to the *bag3^e2/e2^* single mutant*, Tg(cmlc2:tfeb)* single transgenic or WT control fish; n=39-45, log-rank test. **F,** Quantitative RT-PCR analysis of cardiomyopathy molecular markers in *bag3^e2/e2^;Tg(cmlc2:tfeb)* double mutant/transgenic fish compared to single-mutant/transgenic fish and WT control fish. N=3 biological replicates, One-way ANOVA. **G-H,** Western blot (G) and quantification analysis of the LC3 II and ubiquitinated proteins in indicated fish heart treated with or without 50 nM bafilomycin A1 (BafA1) for 4 h. N=4 biological replicates, one-way ANOVA.

### 2. Cardiomyocyte-specific overexpression of *tfeb* d*e*celerated cardiac aging phenotypes in the *bag3* cardiomyopathy model

Upon closer examination of the *bag3^e^*^2^*^/e^*^2^ mutants, we observed a curved body shape in some of *bag3^e2/e2^*mutants at 6 months of age (Supplemental **Fig. 1A**), prompting us to investigate aging-related phenotypes. We detected increased senescence-associated (SA) β-galactosidase activity throughout the body and in the heart at 6 months of age, with more prominent signals in the atrium than that in the ventricle (**Supplemental Fig. 1B**). We next performed immunofluorescent staining of sectioned hearts and observed a significantly increased signal of the cellular senescence marker p16 and DNA damage marker γH2A.X in the cardiomyocytes of *bag3^e2/e2^* mutant fish hearts at 6 months of age (**Fig. 2A-D**). In addition, we carried out quantitative RT-PCR experiments and found significantly elevated transcripts of *p21*, a well-recognized cellular senescence marker in zebrafish ^20^, and senescence-associated secretory phenotype (SASP) markers including *il-1b*, *il-6* and *mmp2* in the *bag3^e2/e2^* mutant fish hearts (**Fig. 2E-2H** and **Supplemental Fig. 1C-F**). Importantly, these aging-associated phenotypic markers were attenuated by the *Tg(cmlc2:tfeb)* transgene **(Fig. 2)**. Together, these results strongly suggest that accelerated cardiac senescence is a pathological event in the *bag3* cardiomyopathy model, which can be ameliorated by cardiomyocyte-specific *tfeb* activation.

**Fig. 2.**
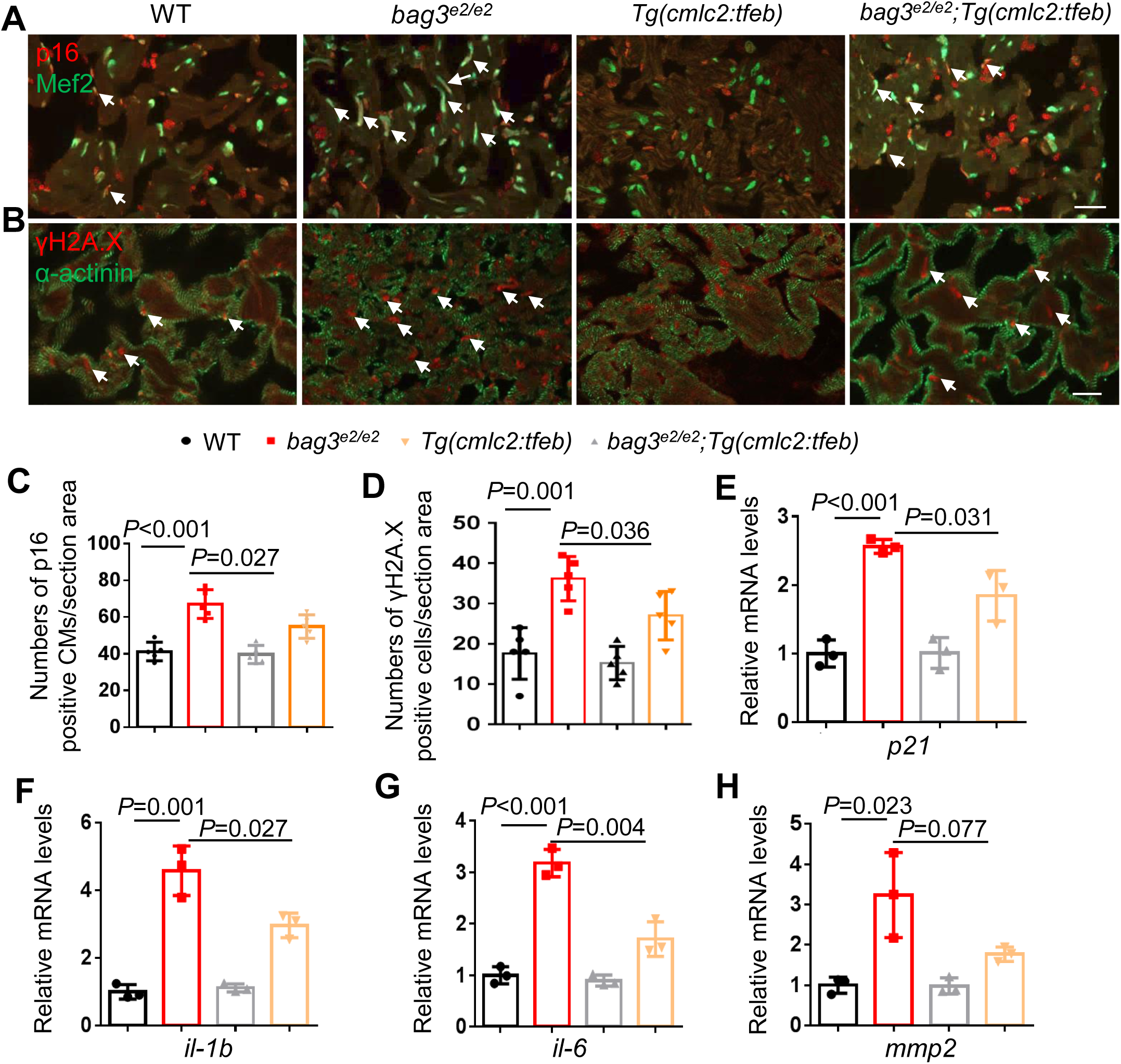
Cardiomyocyte-specific overexpression of *tfeb* decelerated cardiomyocyte senescence. **A-B,** Representative images of cryosectioned heart tissues co-immunostained using either anti-p16 antibody co-stained with anti-Mef2 antibody (A) or anti-γH2A antibody co-stained with anti-α-actinin antibody (B) in the *bag3^e2/e2^;Tg(cmlc2:tfeb)* double- mutant/transgenic fish compared to single-mutant/transgenic fish and WT control fish at 6 months. Scale bars: 20 μm. Arrows point to overlapping signals. **C-D,** Quantification of the numbers of p16/Mef2 and γH2A.X/α-actinin antibodies co-immunostained cells shown in A and B. n=5, One-way ANOVA. **E-H,** Quantitative RT-PCR analysis of cellular senescence marker p21 and senescence-associated secretory phenotype (SASP) markers in *bag3^e2/e2^;Tg(cmlc2:tfeb)* double mutant/transgenic fish compared to single-mutant/transgenic fish and WT control fish. N=3 biological replicates, One-way ANOVA.

### 3. Genetic analysis of shared differentially expressed genes (DEGs) between *Tg(cmlc2:tfeb)* and *mtor^xu0^*^15^*^/+^*identified *fabp7a* as a therapeutic modifier gene for *bag3* cardiomyopathy

To elucidate the molecular mechanisms underlying the cardioprotective effects of *mtor^xu0^*^15^*^/+^*haploinsufficiency mutants and *Tg(cmlc2:tfeb)* transgenes on *bag3* cardiomyopathy, we compared their transcriptomes through RNA-sequencing analysis of whole heart tissues. Principal component analysis revealed that the cardiac transcriptomes of either *mtor^xu0^*^15^*^/+^* haploinsufficiency mutants or *Tg(cmlc2:tfeb)* transgenes formed clusters distinct from the corresponding WT sibling control samples **(Fig. 3A-B**). Using a cut-off of an adjusted *P*-value <0.05 and ≥1.5 fold changes, we identified 1936 differentially expressed genes (DEG) in the *Tg(cmlc2:tfeb)* vs WT, and 65 DEGs in the *mtor^xu0^*^15^*^/+^* vs WT. The top pathways in the *mtor^xu0^*^15^*^/+^*haploinsufficiency mutants include circadian clock, protein folding and macrophage alternative activation signaling pathway. In contrast, the top pathways in the *Tg(cmlc2:tfeb)* transgene lines include cell cycle checkpoints, mitotic prometaphase and activation of the pre-replicative complex. We identified 25 overlapping genes between these two DEG populations, with 15 downregulated and 10 upregulated **(Fig. 3C)**. Intriguingly, among the 15 overlapping downregulated genes, we identified 4 lipodystrophy genes: *bscl2l, fabp7a, cidec* and *edn1* **(Fig. 3D** and **Supplemental Table 1).** We chose to test potential modifying effects of these 4 lipodystrophy genes on the *bag3* cardiomyopathy model using an F0- based genetic assay ^21^. We designed microhomology-mediated end joining (MMEJ)- inducing single guide RNAs (sgRNAs) and tested their knockout efficiency via injecting into wildtype embryos. After confirming high knockout efficiency, i.e. ranging from 76% to 90% **(Fig. 3E)**, we injected each of the 4 sgRNAs into the offsprings of *bag3^e2/+^* incrosses. We found that only the sgRNA targeting the first exon of *fabp7a* gene *(fabp7a^MJ-e1^*) restored the cardiac dysfunction in the *bag3* cardiomyopathy model at 6 months of age **(Fig. 3F)**. This suggests that *fabp7a* is a potential therapeutic modifier gene for *bag3* cardiomyopathy.

**Fig. 3.**
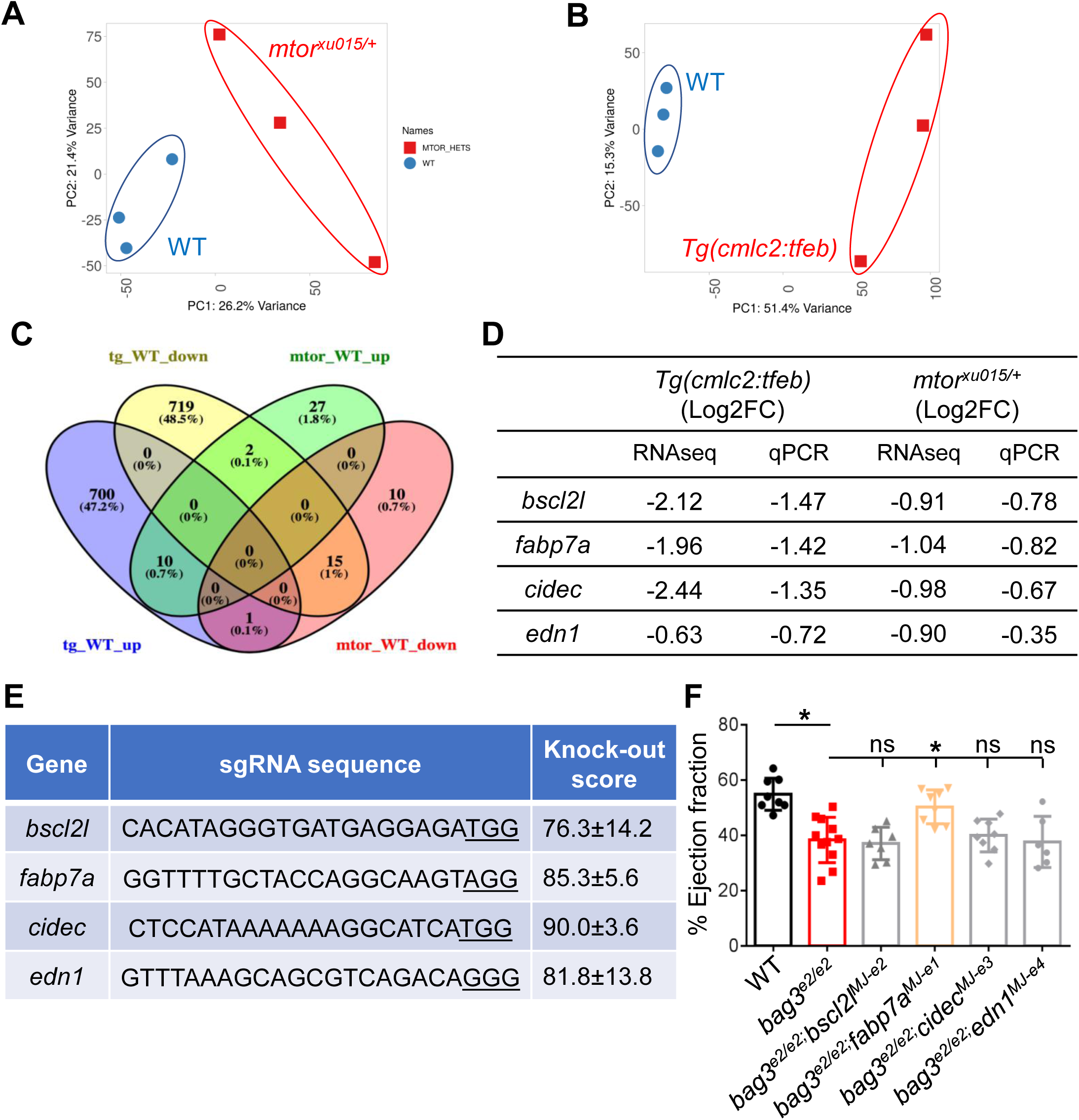
Combination of RNA-seq with an F0-based genetic screen identified *fabp7a* as a therapeutic modifier gene for the *bag3* cardiomyopathy model. **A-B,** Principal Component Analysis (PCA) of RNA-seq-based expression data from the *mtor^xu0^*^15^*^/+^*mutant (A) or *Tg(cmlc2:tfeb)* (B) transgenic fish hearts compared to WT controls. C, Comparison of cardiac transcriptome between *mtor^xu0^*^15^*^/+^* mutant and *Tg(cmlc2:tfeb)* transgenic fish hearts identified 10 upregulated and 15 downregulated genes as common DEGs. **D,** Quantitative RT-PCR validated the expression of 4 lipodystrophic DE genes that were downregulated in both the *mtor^xu0^*^15^*^/+^* mutant and the *Tg(cmlc2:tfeb)* transgenic hearts. **E,** List of single guide RNA (sgRNA) sequences for the 4 lipodystrophic DE genes and knockout scores detected from the adult fish injected with sgRNAs in F0 generation. **F,** Injection of *fabp7a* microhomology-mediated end joining (MMEJ)-inducing sgRNA, but not the other 3 individual MMEJ sgRNAs, exerts cardioprotective effect on *bag3^e2/e2^* cardiomyopathy in F0 adult fish.

Next, we performed a Western blot analysis and detected significantly elevated expression of the Fabp7a protein in the *bag3* cardiomyopathy model compared to WT control fish hearts (**Fig. 4A**). To confirm the salutary modifying effect of *fabp7a* inhibition on *bag3* cardiomyopathy, we generated a stable *fabp7a* mutant that harbors an 8- nucleotide deletion (**Fig. 4B**). At the protein level, the expression level of Fabp7a was reduced to 57% in the *fabp7a^e1/+^*heterozygous mutant and 3% in the *fabp7a^e1/e1^* homozygous mutant fish compared to that in WT control (**Fig. 4C**). We found that cardiac dysfunction in the *bag3^e2/e2^* mutant fish at 6 months of age could be partially rescued in the *bag3^e2/e2^;fabp7a^e1/+^*double-mutant fish (**Fig. 4D**). Both sarcomeric damage and mitochondrial swelling phenotypes were largely restored in the *bag3^e2/e2^;fabp7a^e1/+^* double-mutant fish (**Fig. 4E**). At the molecular level, re-activation of fetal gene programs detected in the *bag3^e2/e2^* mutant hearts was partially inhibited in the *bag3^e2/e2^;fabp7a^e1/+^* double-mutant hearts **(Fig. 4F)**. Collectively, these data confirm *fabp7a* as a modifier gene for *bag3* cardiomyopathy that can be modulated to exert cardioprotective effects.

**Fig. 4.**
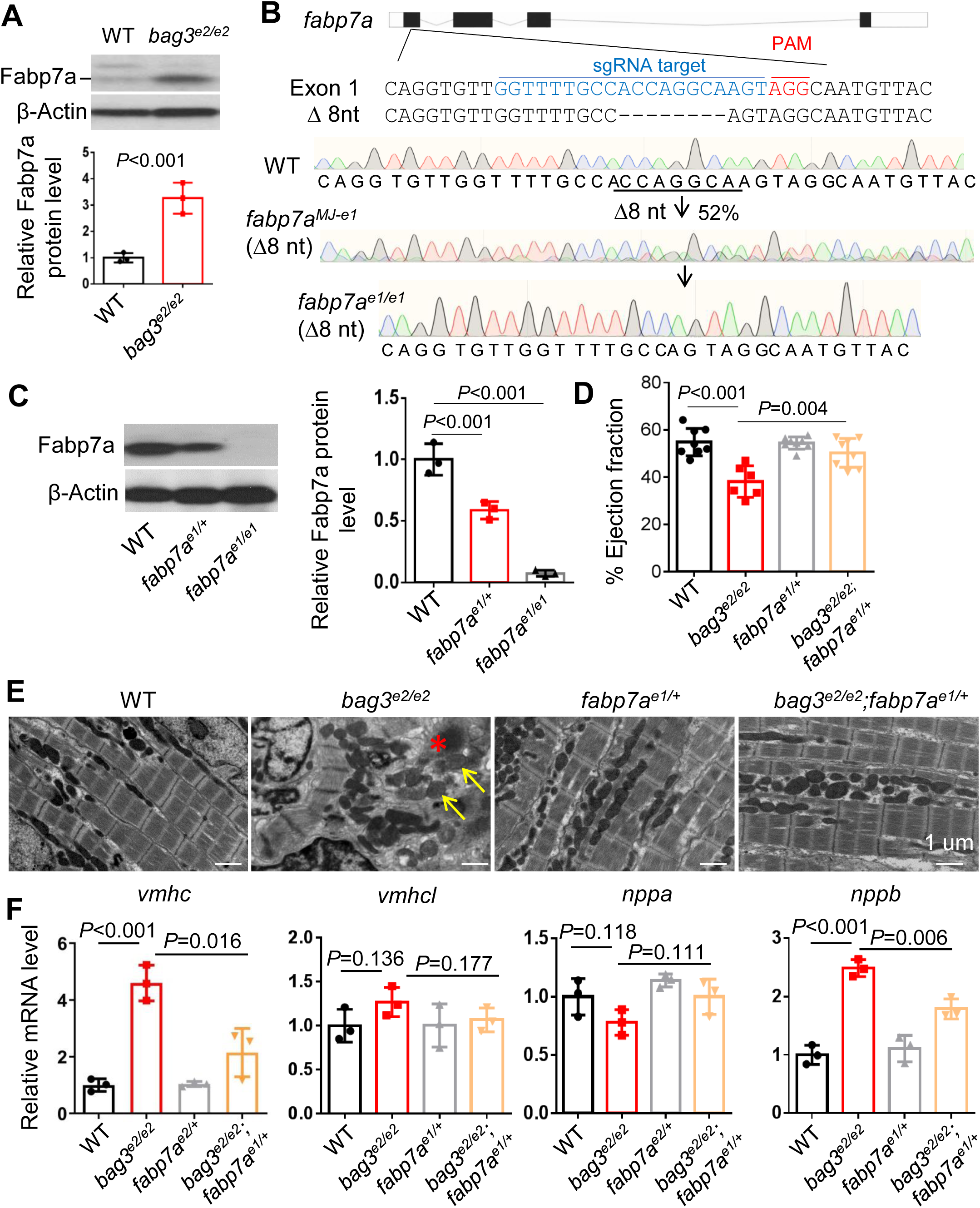
The therapeutic modifying effects of *fabp7a* inhibition was validated in the stable F1 haploinsufficiency mutant. **A,** Western blot and quantification analysis of the Fabp7a protein levels in heart lysates of *bag3^e2/e2^* mutant compared to WT control fish hearts at 6 months. **B,** Schematic and chromatographs illustrating the genetic lesion of 8-nucleotides generated by injection of an MMEJ inducing *fabp7a* sgRNA targeting sequences within the first exon. Dashed lines indicate an 8-nucleotide-long deletion. **C,** Western blot and quantification analysis of the Fabp7a protein levels in the *fabp7a* stable heterozygous (*fabp7a^e1/+^*) and homozygous (*fabp7a^e1/e1^*) mutant fish compared to WT controls. n=3, one-way ANOVA. **D,** Quantification of cardiac function. Ejection fraction (EF) (in %) measured by echocardiography in *bag3^e2/e2^;fabp7a^e1/+^* double-mutant fish compared to single-mutant and WT control fish at 6 months; n=6-8, one-way ANOVA. **E,** Confirmative TEM images of *bag3^e2/e2^;fabp7a^e1/+^* double-mutant fish hearts compared to *bag3^e2/e2^* and *fabp7a^e1/+^* single mutant and WT controls at 6 months. The asterisk indicates Z-disc aggregation. Arrows point to mitochondrial rounding and swelling. Scale bar: 2 μm. **F,** Quantitative RT-PCR analysis of cardiomyopathy molecular markers in *bag3^e2/e2^;fabp7a^e1/+^* double-mutant fish compared to single-mutant and WT control fish at 6 months. N=3 biological replicates, one-way ANOVA.

### 4. *fabp7a* inhibition restored dysregulated proteostasis and decelerated cardiac aging phenotypes in the *bag3* cardiomyopathy model

Because dysregulated proteostasis is a major pathological feature of *bag3* cardiomyopathy, we investigated whether *fabp7*a inhibition could repair proteostasis dysregulation. Indeed, our Western blot results showed that the proteostasis defects in the *bag3^e2/e2^* mutant hearts, as indicated by the aberrant response of LC3 II to BafA1 treatment and elevated ubiquitinated protein aggregation, were either rescued or partially rescued by the *fabp7a^e1/+^* haploinsufficiency mutation **(Fig.5A-B)**. Given that the *bag3* cardiomyopathy model exhibited accelerated cardiac aging phenotypes, we then asked whether *fabp7*a inhibition affects cardiomyocyte senescence through immunostaining assays. We found that the elevated expression of cellular senescence marker p16 and DNA damage marker γH2A.X in the *bag3^e2/e2^* mutant hearts were significantly ameliorated in the *bag3^e2/e2^;fabp7a^e1/+^* double-mutant hearts (**Fig. 5C-E**). In addition, at the transcript level, the elevated expression of cellular senescence marker *p21* and SASP genes including *il-1b* and *il-6,* but not *mmp2,* in the *bag3^e2/e2^*mutant hearts were also partially inhibited in the *bag3^e2/e2^;fabp7a^e1/+^* double mutant hearts (**Fig. 5F-I).** Together, these results suggest that *fabp7a* inhibition can repair dysregulated proteostasis and rejuvenate cardiomyocytes in the *bag3* cardiomyopathy model.

**Fig. 5.**
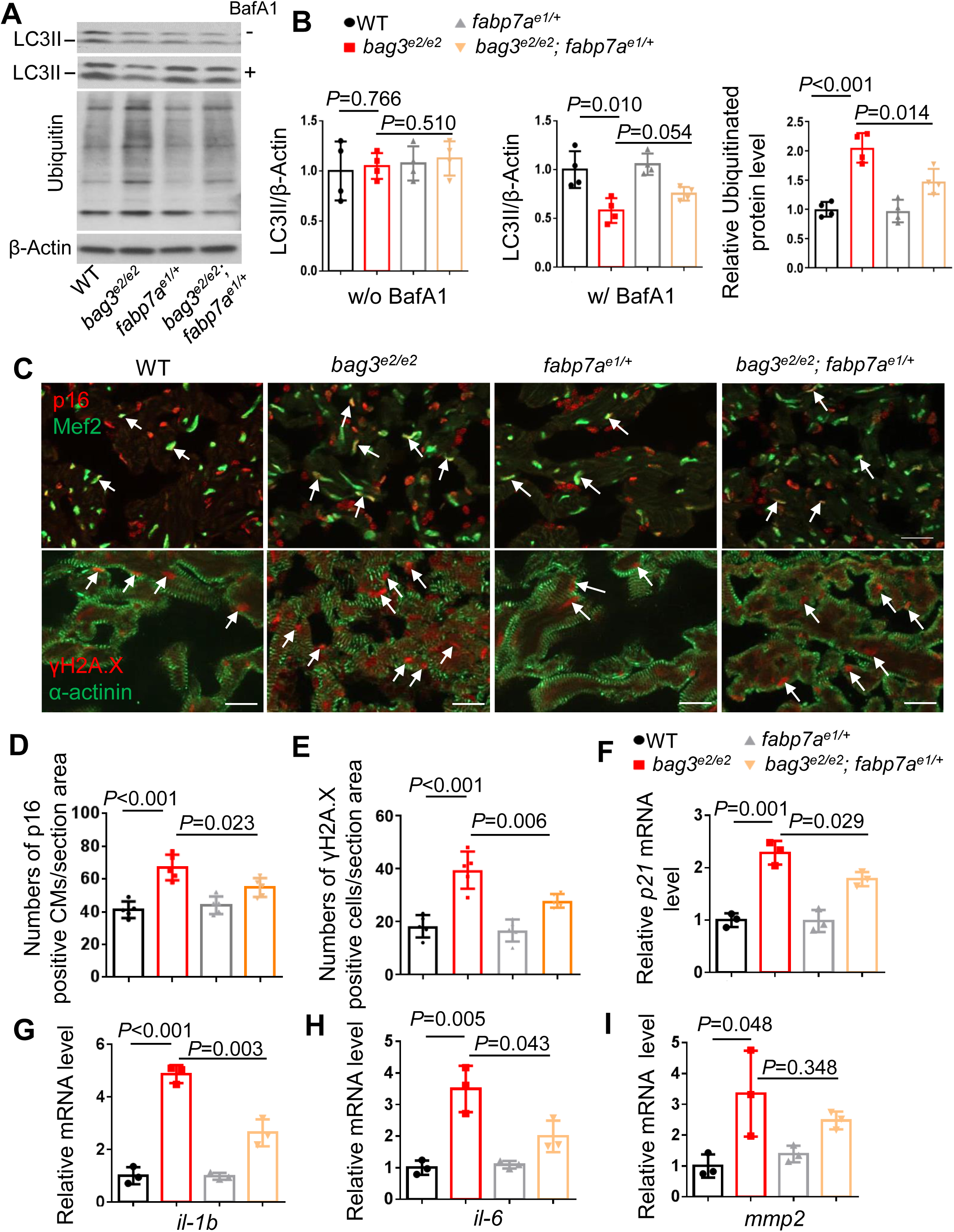
fabp7a inhibition is sufficient to restore impaired protein homeostasis and decelerate cardiomyocyte senescence in the *bag3* cardiomyopathy model. **A-B,** Western blot (A) and quantification analysis (B) of LC3II and ubiquitinated protein levels in heart lysates of *bag3^e2/e2^;fabp7a^e1/+^* double-mutant, *bag3^e2/e2^, fabp7a^e1/+^* single mutant or WT control fish hearts at 6 months treated with or without 50 nM BafA1 for 4 h. n=4, one-way ANOVA. **C,** Representative images of immunostaining using either anti-p16 antibody co-stained with anti-Mef2 antibody (A) or anti-γH2A.X antibody co-stained with anti-α-actinin antibody (B) in *bag3^e2/e2^;fabp7a^e1/+^* double-mutant, *bag3^e2/e2^, fabp7a^e1/+^* single mutant or WT control fish hearts at 6 months. Arrows point to overlapping signals. Scale bars: 20 μm. **D-E,** Quantification of the numbers of p16/Mef2 (D) and γH2A.X/α-actinin (E) antibodies co-immunostained cells shown in C. n=5, One-way ANOVA. **F-I,** Quantitative RT-PCR analysis of cellular senescence marker p21 and SASP markers in *bag3^e2/e2^;fabp7a^e1/+^*double-mutant compared to single-mutant fish and WT control fish hearts at 6 months. N=3 biological replicates, One-way ANOVA.

### 5. Cardiomyocyte-specific overexpression of *fabp7a* caused cardiomyopathy accompanied by dysregulated proteostasis and accelerated cardiac aging phenotypes

To further elucidate the functions of *fabp7a* in cardiomyopathy, we generated a conditional overexpression line named *Tg(*β*-actin2:loxP-mCherry-loxP-fabp7a-cerulean),* or *Tg(fabp7a-OE)* for simplicity in adult zebrafish **(Fig. 6A).** After crossing the *Tg(fabp7a-OE)* fish with the *Tg(cmlc2-creERT2)* line that drives cardiomyocyte-specific Cre expression upon 4-Hydroxytamoxifen (4-HT) induction, we observed an obvious fluorescence switch from mCherry to cerulean in the hearts of the *Tg(fabp7a- OE)*;*Tg(cmlc2-creER)* double transgenic fish at one-week post-4-HT treatment. The ectopic expression of the Fabp7a-cerulean fusion protein in the double transgenic fish hearts was confirmed by Western blot analysis **(Fig. 6B-C).** Next, we carried out high frequency echocardiography analysis and detected a significantly reduced cardiac function in the *Tg(fabp7a-OE);Tg(cmlc2-creER)* double transgenic fish at three months post-4-HT treatment **(Fig. 6D)**. Transmission electron microscopy (TEM) analysis confirmed impaired sarcomere structure and mitochondrial swelling defects **(Fig. 6E)**. At the molecular level, aberrant expression of cardiomyopathy markers including *vmhc*, *vmhcl* and *nppb* were detected, further supporting the cardiomyopathy-like phenotypes in the T*g(fabp7a-OE);Tg(cmlc2-creER)* double transgenic fish **(Fig. 6F)**. We next conducted Western blot analysis and detected dysregulated protein homeostasis in the *Tg(fabp7a-OE);Tg(cmlc2-creER)* double transgenic fish at three months post-4HT treatment, as indicated by stalled response of LC3II to BafA1 and elevated ubiquintinated protein aggregation **(Fig. 7A-C)**. Additionally, we examined cardiac aging indices and detected elevated staining for p16 and γH2A.X in cardiomyocyte **(Fig. 7D- E)**, as well as upregulated mRNA expression of *p21* and SASP genes including *il-1b* and *il-6* in the *Tg(fabp7a-OE);Tg(cmlc2-creER)* double transgenic fish at 3 months post- 4HT treatment **(Fig. 7F)**. Collectively, these results suggest that cardiomyocyte-specific overexpression of *fabp7a* caused cardiomyopathy accompanied by dysregulated proteostasis and accelerated cardiac aging phenotypes.

**Fig. 6.**
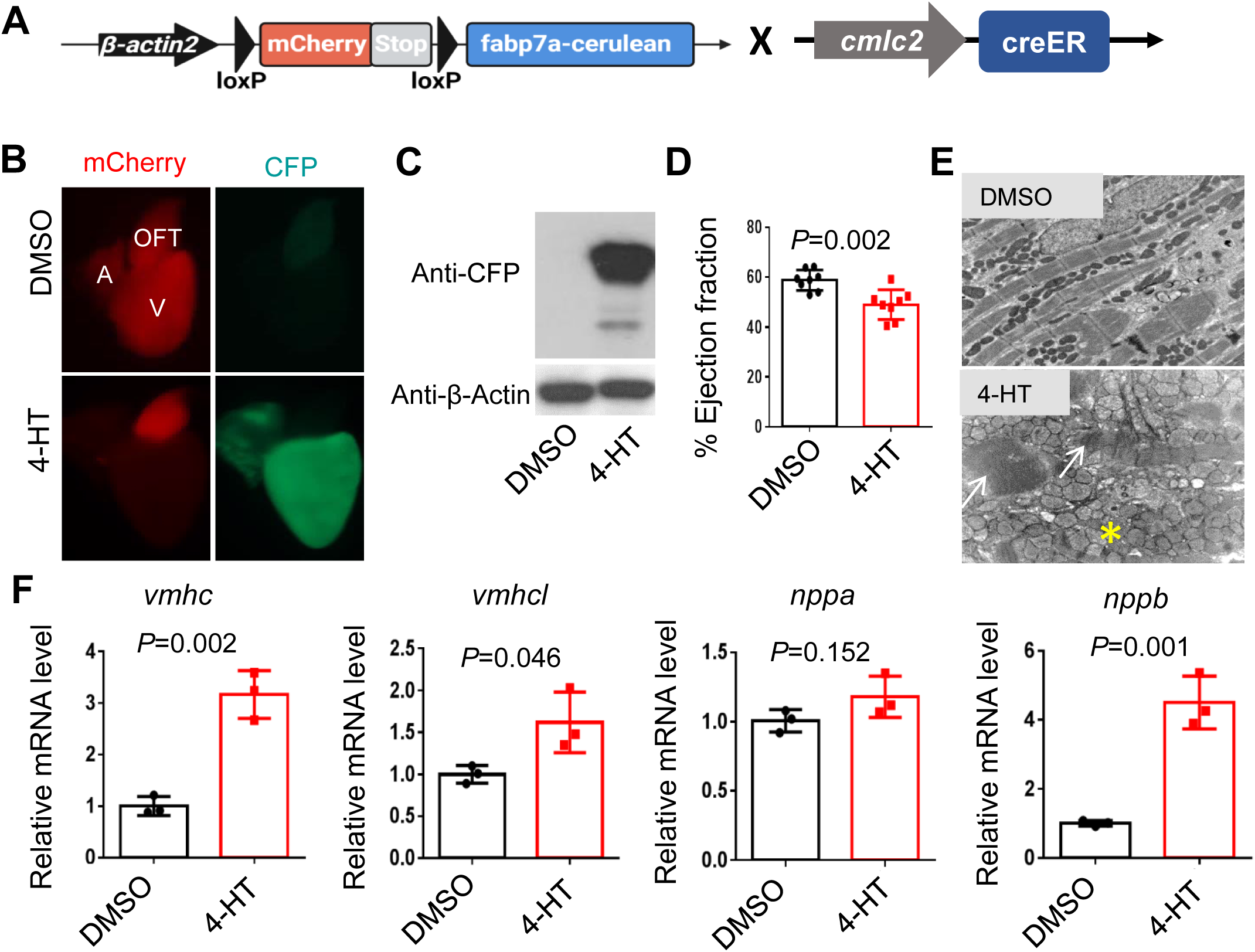
Cardiomyocyte-specific overexpression of *fabp7a* gene results in cardiac dysfunction. **A,** Schematic constructs and design to generate a transgenic line with cardiomyocyte-specific overexpression of *fabp7a* in zebrafish. **B-C**, Cardiomyocyte-specific overexpression of Fabp7a was indicated by fluorescence switch from mCherry to cerulean fluorescent protein (CFP) (B) and by Western blot analysis (C) at one-week post-4-Hydroxytamoxifen (4-HT) induction. **D-E**, Cardiomyocyte-specific overexpression of Fabp7a led to cardiac function decline (D), impaired sarcomere (arrows) and swollen mitochondria (asterisk) (E) at 3 months post-4-HT induction. **F,** Quantitative RT-PCR analysis of cardiomyopathy molecular markers in the cardiomyocyte-specific overexpression of Fabp7a fish at 3 months post-4-HT induction. N=3 biological replicates, one-way ANOVA.

**Fig. 7.**
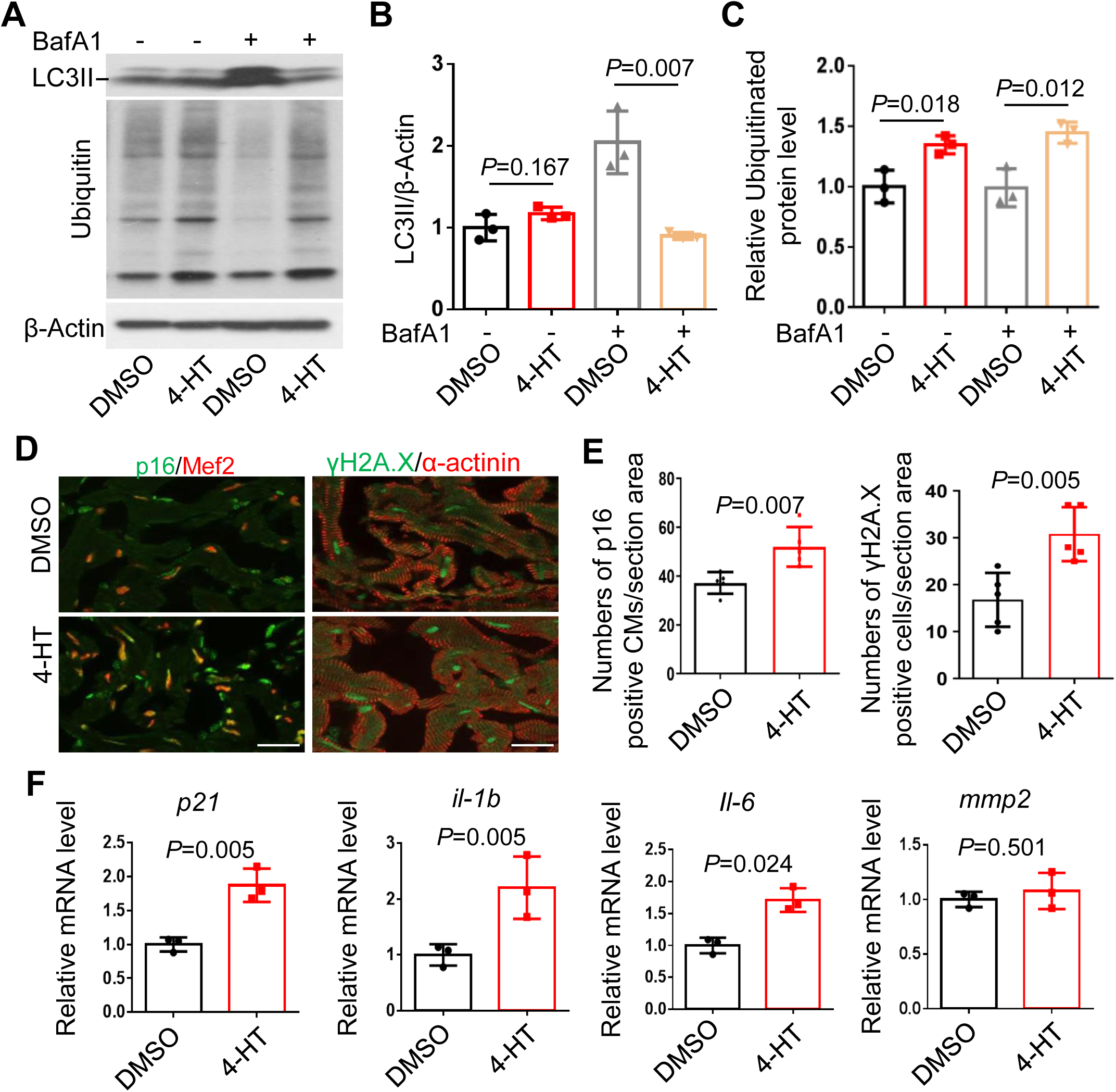
Cardiomyocyte-specific overexpression of *fabp7a* gene caused impaired protein homeostasis and accelerated cellular senescence. **A-B,** Western blot (A) and quantification analysis of the LC3 II (B) and ubiquitinated proteins (B) levels indicated fish heart treated with or without 50 nM BafA1 for 4 h. N=3 biological replicates, one-way ANOVA. **D,** Representative images of immunofluorescence using anti-p16 antibody co-stained with anti-Mef2 antibody or anti-γH2A.X antibody co-stained with anti-α-actinin antibody in the cardiomyocyte-specific overexpression of Fabp7a fish at 3 months post-4-HT induction. Scale bars: 20 μm. **E,** Quantification of the numbers of anti-p16/anti-Mef2 and anti-γH2A.X/anti-α-actinin co-stained cells shown in G. N=5, One-way ANOVA. F, Quantitative RT-PCR analysis of cellular senescence marker p21 and SASP markers in the cardiomyocyte-specific overexpression of Fabp7a fish at 3 months post-4-HT induction. N=3 biological replicates, one-way ANOVA.

## Discussion

### 1. Accelerated cardiac aging and increased Fabp7a expression are two pathological events in the *bag3* cardiomyopathy model

Prior to the present study, it was well established that dysregulated proteostasis is a primary pathological event in *bag3* cardiomyopathy, significantly contributing to its pathogenesis ^22, 23^. In this manuscript, we further unveiled that accelerated cardiac senescence also occurs in the *bag3* cardiomyopathy model, as has been reported previously in the AIC model. Our data suggest that accelerated cardiac senescence may be a common pathological feature across different cardiomyopathy etiology. Given that dysregulated proteostasis is one of the twelve hallmarks of normative aging ^12^, it is possible that accelerated cardiac senescence is either correlated with, or even results from dysregulated proteostasis in the heart. More detailed mechanistic studies are warranted to explore these relationships in detail.

In addition, our genetic screen identified upregulated Fabp7 expression as an important molecular event in *bag3* cardiomyopathy associated with both dysregulated proteostasis and accelerated cardiac senescence (**Fig. 8**). Phylogenetic analysis indicated that human FABP7 is the closest family member to FABP3, a cardiac-type FABP, sharing the highest amino acid identity with it compared to other FABP members^14^. Contrary to the previous understanding that FABP7 is a brain-type FABP primarily functions in the brain, our data reveal novel cardiac expression and functions of Fabp7. Transgenic studies demonstrated that cardiomyocyte-specific overexpression of *fabp7a* is sufficient to drive proteostasis abnormality and accelerate cardiac senescence.

**Fig. 8.**
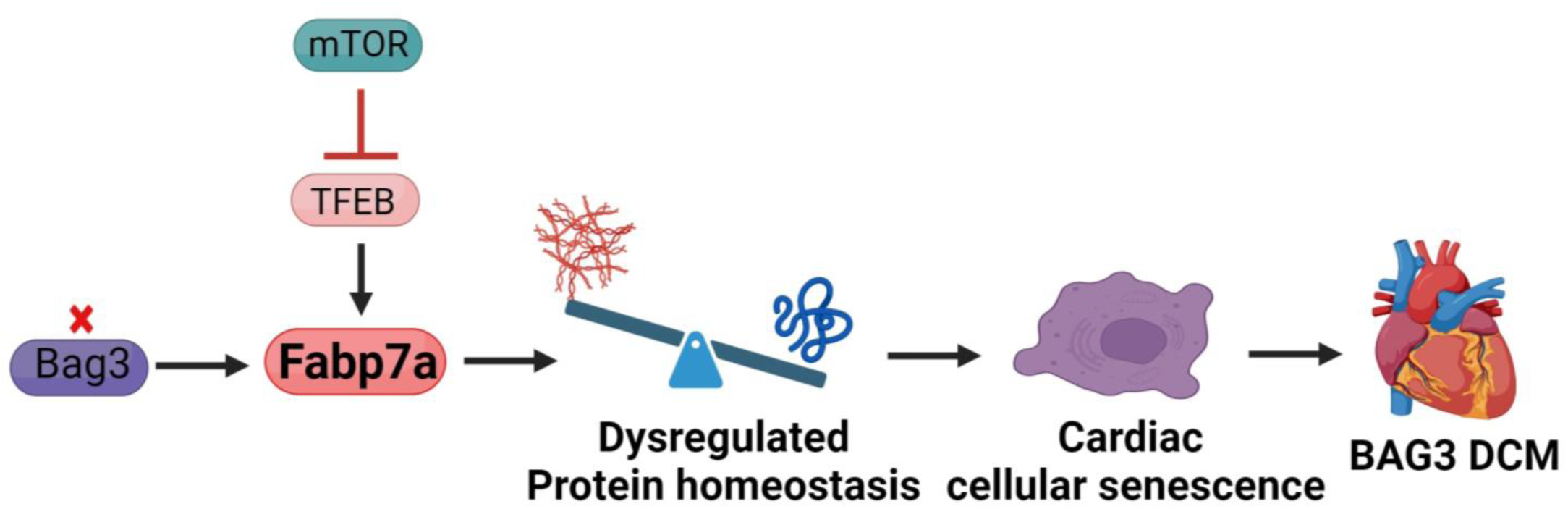
Schematics for the main discoveries. The mTOR-Tfeb-Fabp7a signaling axis regulated protein homeostasis and cardiac aging in *bag3* cardiomyopathy.

Because Fabp7 is primarily involved in lipid homeostasis, our data suggest an intrinsic link between lipid metabolism and proteostasis that could be important for pathogenesis of *bag3* cardiomyopathy. Further experimental evidence is needed to test this hypothesis and elucidate the underlying mechanisms.

### 2. The mTOR-TFEB-Fabp7 signaling axis can be harnessed for therapeutic benefits in *bag3* cardiomyopathy

We have previously shown that inhibiting mTOR signaling through the *mtor^xu0^*^15^*^/+^*haploinsufficiency mutant exerted cardioprotective effects on *bag3* cardiomyopathy ^6^. Here, we further reported that cardiomyocyte-specific overexpression of the zebrafish *tfeb* ortholog, a direct downstream target of mTOR signaling pathway, ameliorated dilated cardiac structure remodeling, rescued the cardiac function decline, and prolonged survival in the *bag3* cardiomyopathy model. These results thus suggest that activation of *tfeb* may confer the cardioprotective effect of mTOR inhibition on *bag3* cardiomyopathy. Consistent with its well-recognized role in proteostasis ^24^, activation of *tfeb* reduced the accumulation of ubiquitinated protein aggregation in the *bag3* cardiomyopathy hearts and attenuated cellular senescence.

Downstream of the mTOR-Tfeb axis, *fabp7a* is a candidate therapeutic target gene for *bag3* cardiomyopathy. While expression of *fabp7a* mRNA was downregulated in the hearts of the *mtor^xu0^*^15^*^/+^* haploinsufficiency mutants and *Tg(cmlc2:tfeb)* transgenes, it was aberrantly elevated in the *bag3* cardiomyopathy model. Haploinsufficiency mutation in *fabp7a* mostly rescued the cardiomyopathy phenotypes and inhibited accelerated cardiac aging indices in the *bag3* cardiomyopathy model. Conversely, cardiomyocyte-specific overexpression of *fabp7a* led to elevated expression of cellular senescence markers and caused cardiomyopathy. Together, these genetic studies suggest an mTOR-Tfeb-Fabp7a signaling axis that can be harnessed to exert therapeutic benefits in *bag3* cardiomyopathy (**Fig. 8**). Mechanistically, this signaling axis can be manipulated to repair dysregulated proteostasis and attenuate accelerated cardiac senescence. While *mtor* and *fabp7a* need to be inhibited, *tfeb* should be activated in the cardiomyocytes to exert therapeutic benefits. Future studies are warranted to elucidate how Tfeb negatively regulates Fabp7a expression. Given that TFEB has been reported to bind to the promoter regions of other human FABP family members such as FABP3, FABP4 and FABP9 to directly regulate their transcriptional expression ^17^, it is reasonable to speculate that Tfeb might directly bind to the promoter of Fabp7a to control its transcriptional activity. Consistent with this concept, transcription factor binding motif analysis revealed four potential Tfeb binding sites in the upstream promoter region of *fabp7a* (Supplemental Fig. 3).

### 3. Experimentally testing DEGs using an F0-based genetic assay is an effective strategy for discovering genetic modifiers of cardiomyopathies

The identification of *fabp7a* as a therapeutic modifier gene downstream of the mTOR-Tfeb signaling axis underscored the power of zebrafish genetics, highlighting an F0-based genetic assay as a rapid strategy for discovering novel genetic modifiers of inherited cardiomyopathy. As demonstrated in our recent finding ^21^, designing a microhomology-mediated end joining (MMEJ) sequence target for single guide RNAs allows for obtaining high knockout scores with. knockout scores ranging from 76% to 90% for each of the individual candidate genes in this study. These relatively high knockout scores allow us to test the modifying effects of loss-of-function mutation in candidate genes on *bag3* cardiomyopathy within the F0 generation, significantly bypassing the lengthy multi-generational genetic crosses required for assessing the effects of candidate modifying genes in a disease setting. In the era of genomics, where a vast amount of candidate genes is identified through sequencing or transcriptome analysis, this F0-based genetic screening approach provides a rapid method to identify genetic modifiers. This efficient strategy for *bag3* cardiomyopathy is anticipated to be extendable to other types of cardiomyopathies or any human diseases that can be modeled in zebrafish.

## Methods

### Sex as a biological variable

Our study examined male and female animals, and similar findings are reported for both sexes.

### Animals

Zebrafish *(Danio rerio)* were maintained under a 14-hour light–10-hour dark cycle at

28.5°C and handled with care. The animal study protocols were approved by the Mayo Clinic Institutional Animal Care and Use Committee (A3531 for zebrafish). All animal study procedures were performed in accordance with the Guide for the Care and Use of Laboratory Animals published by the US National Institutes of Health (NIH Publication No. 85-23, revised 1996).

### Measurement of the ventricular surface area to body weight ratio

The ventricular surface area (VSA) to body weight (BW) ratio was measured according to a previously published method ^25^. To measure the VSA, the ventricles of individual zebrafish were dissected and imaged next to a millimeter ruler under a Leica MZ FLI III microscope. The largest projection of a ventricle was outlined using ImageJ software. To measure body weight, the fish were anesthetized in 0.16 mg/mL tricaine solution, semi-dried on a paper towel, and weighted on a scale. The VSA/BW index was then calculated by the largest projection area of ventricle (in mm^2^) divided by the body weight (in gram) ^26^.

### In vivo echocardiography for adult zebrafish hearts

The Vevo 3100 high-frequency imaging system equipped with a 50 MHz (MX700) linear array transducer (FUJIFILM VisualSonics Inc) was used to measure cardiac function indices in adult zebrafish according to a reported protocol ^26^. Briefly, acoustic gel (Aquasonic® 100, Parker Laboratories, Inc) was applied to the surface of the transducer to ensure adequate coupling with the tissue interface. Adult zebrafish at appropriate ages were anesthetized in 0.16 mg/mL tricaine for 3 minutes and placed ventral side up into a sponge. The MX700 transducer was placed above the zebrafish to provide a sagittal imaging plane of the heart. B-mode images were acquired with an imaging field view of 9.00 mm in the axial direction and 5.73 mm in the lateral direction, a frame rate of 123 Hz, with medium persistence and a transmit focus at the center of the heart. Image quantification was performed using the VevoLAB workstation. For each index on individual fish, measurements wereconducted on 3 independent cardiac cycles to obtain average values.

### Transmission electron microscopy

For the TEM study, adult zebrafish hearts were dissected and immediately fixed in Trump’s solution (4% paraformaldehyde and 1% glutaraldehyde in 0.1 M phosphate buffer [pH, 7.2]) at room temperature (RT) for 1 hour, followed by overnight incubation at 4°C. The fixed samples were subsequently processed and imaged at the Mayo Clinic Electron Microscopy Core Facility using a Philips CM10 transmission electron microscope.

### Quantitative real-time PCR

Total RNA was extracted from an individual adult fish ventricle using Trizol reagent (ThermoFisher Scientific) following the manufacturer’s instruction. Approximately 100 ng total RNA was used for reverse transcription (RT) and cDNA synthesis using Superscript III First-Strand Synthesis System (ThermoFisher Scientific). Real-time quantitative PCR was performed in 96-well optical plates (ThermoFisher Scientific) using an Applied Biosystem VAii 7 System (ThermoFisher Scientific). Gene expression levels were normalized using the expression level of either glyceraldehyde 3-phosphate dehydrogenase (*gapdh) or actin, beta 2 (actb2)* by –ΔΔCt (cycle threshold) values. Primer information for qRT-PCR is listed as follows:

*nppa*-F: 5’-GATGTACAAGCGCACACGTT-3’, *nppa-*R: 5’- TCTGATGCCTCTTCTGTTGC-3’; *nppb*-F: 5’-CATGGGTGTTTTAAAGTTTCTCC-3’, *nppb*-R, 5’-CTTCAATATTTGCCGCCTTTAC-3’; *vmhc*-F, 5’- TCAGATGGCAGAGTTTGGAG5-3’,*vmhc*-R, 5’-GCTTCCTTTACAGTTACAGTCTTTC5- 3’; *vmhcl*-F: 5’- GCGATGCTGAAATGTCTGTT-3’,*vmhcl*-R, 5’- CAGTCACAGTCTTGCCTCCT-3’; *p21-F,* 5’-AGGAA AAG CAGCAG AAACG-3’, *p21-R*, 5’-TGTTG GTCTGT TTG CGCTT-3’; *il-1b*-F, 5’-CTGGAGATGTGGACTTCGCA-3’ *il-1b*-R, 5’--TCACGCTCTTGGATGACGTT-3’; *il-6*-F, 5’- CAGAGACGAGCAGTTTGAGAGA-3’, *il-6*-R, 5’-TCAGGACGCTGTAGATTCGC5-3’; *mmp2-F,* 5’-AGCTTTGACGATGACCGCAAATGG-3’, *mmp2-R,* 5’- GCCAATGGCTTGTCTGTTGGTTCT-3’; β*-actin*-F, 5’-TTCACCACCACAGCCGAAAGA- 3’, β*-actin*-R, 5’-TACCGCAA GATTCCATACCCA-3’.

### Antibody immunostaining

Heart samples dissected from adult zebrafish were embedded in a tissue freezing medium and sectioned at 8 μm using a cryostat (Leica CM3050 S). Sections were air dried for 30 minutes (mins) atRT and fixed with 4% PBS-buffered paraformaldehyde (PFA) for 7 mins, permeabilized with 0.1% Triton X-100 in PBD (1X PBS, 1% BSA, 1% DMSO) for 45 mins, blocked with 2% sheep serum/PBD for 25 mins, and incubated with primary antibodies overnight at 4℃. Primary antibody-stained sections were then washed in PBD for three times and incubated with secondary antibodies (Alexa Fluor anti-Rabbit 488, ThermoFisher Scientific, catalog #A11008; Alexa Fluor anti-Mouse 568, ThermoFisher Scientific, catalog #A11001) at RT for 1 hour, washed with PBD for three times and transferred to a slide with a mounting medium with DAPI (Vector, H-1200). Stained sections were imaged with a Zeiss Axioplan 2 microscope equipped with ApoTome and AxioVision software (Carl Zeiss). The following primary antibodies were used: anti-MEF2(A+C) (1:300, Abcam, catalog #197070), anti-p16INK4a (1:100, Santa Cruz, catalog #sc-1661), anti-γH2A.X (phosphor Ser139) (1:200, GeneTex, catalog #GTX127342), anti-α-actinin (1:300, Sigma, catalog #A7811), anti-Fabp7 (1:100, GeneTex, catalog #GTX121467).

### Western blotting

Either pooled zebrafish embryos at 3 days post-fertilization or individually dissected adult zebrafish heart ventricles were transferred to the RIPA lysis buffer (Sigma-Aldrich) supplemented with complete protease inhibitor cocktail (Roche) and homogenized using a Bullet Blender tissue homogenizer (Next Advance, Inc). The resultant protein lysates were subjected to western blotting using a standard protocol. The following primary antibodies were used: anti-γH2A.X (phosphor Ser139) (1:1,000, GeneTex, catalog #GTX127342), anti-H3k9me3 (1:5,000, Diagenode, catalog #C15410056), anti-LC3 (1:2,000, Cell Signaling Technology, catalog #12741), anti-β-actin (1:8,000, Santa Cruz Biotechnology Inc., catalog #sc-1615), anti-Ubiquitin (1:1,000, ThermoFisher Scientific, catalog #PA5-17067; or 1:1,000, Cell Signaling Technology, catalog #3936), anti-Fabp7 (1:1,000, GeneTex, catalog #GTX121467).

### RNA-seq analysis

RNA-seq data acquisition and analysis were performed according to previous published protocol ^6, 26^. Briefly, total RNA was extracted from dissected ventricular tissue of 6-month-old *mtor^xu0^*^15^*^/+^* mutant, *Tg(cmlc2:tfeb)* transgenic fish and corresponding WT siblings. Five ventricles were pooled as one sample and three biological replicates for each genotype were sequenced using the HiSeq 2000 platform (Illumina) with a 50-bp paired-end sequencing protocol in the Mayo Clinic DNA Sequencing Core Facility. Raw RNA-seq reads for each sample were originally aligned with TopHat (Version 2.1.1) to the zebrafish genome assembly (Zv9) using the Ensembl annotation Zv9 (Danio_rerio.Zv9.79.gtf) and later with the updated genome assembly GRCz11 (Genome assembly GRCz11). Gene expression was quantified using Cufflinks (Version 2.2.1). Differential gene expression across different groups was determined based on fold change and a false discovery rate of less than 0.05 according to the Cuffdiff script from Cufflinks. Unsupervised hierarchical clustering was performed using Pearson correlation and scaled based on the fragments per kilobase of transcript per million mapped reads value with the pheatmap R package (https://github.com/raivokolde/pheatmap). Both the primary RNA-seq raw and processed datasets have been deposited in GEO under the accession number GSE269725.

### MMEJ-based single guide RNA design and F0 based genetic testing

MMEJ-based single guide RNA design and F0 injection were performed according to a recently published protocol ^21^. Briefly, targeted exon sequences of genes were uploaded to an online algorithm, MENTHU (http://genesculpt.org/menthu/). Target guide RNA sequences with high scores were selected from predicted MMEJ loci. Single guide RNAs (sgRNAs) with appropriate chemical modifications were synthesized and obtained from Synthego (Synthego Corporation). sgRNAs were dissolved in nuclease-free duplex buffer (Integrated DNA Technologies, 11-01-03-01) and diluted to 5 µM as working concentrations. The sgRNA-Cas9 protein (sgRNP) complex was then assembled and injected into one-cell staged zebrafish embryos to obtain MMEJ-injected F0 embryos. Individual MMEJ sgRNP-injected embryos were harvested and subjected to knockdown efficiency measurement and calculation. Sequences were obtained via Sanger sequencing at Genewiz (https://clims4.genewiz.com/CustomerHome/Index), and the knockdown scores were obtained using the R code-based open access software Inference of CRISPR Edits (ICE) v2 CRISPR Analysis Tool (https://www.synthego.com/products/bioinformatics/crispr-analysis) ^27^. Primer information for assessing knockdown efficiency is listed as follows: bscl2-MJ-F1, 5’-TGAACAGAAATGGAGCGTG-3’, bscl2-MJ-R1, AATCAAGAATGGTCACCTGATG-3’; fabp7a-MJ-F1, 5’-GCATGTGTAAGGTGCAGTAG- 3’, fabp7a-MJ-R1, 5’-GTGGTTTCATCAAACTCCTCTC-3’; cidec-MJ-F1, 5’- ATGTGCGTTGTGTGTCAC-3’, cidec-MJ-R1, 5’-TGTTTGCCGGGTTCTTTC-3’; edn1- MJ-F1, AGCTGTCATTGCATGACTTG-3’, edn1-MJ-R1, 5’- CACTGTCTCTGTGGTTTGTC-3’.

### Generation of cardiomyocyte-specific overexpression of *fabp7a* transgenic line

The *Tg(*β*actin2:loxP-mCherry-stop-loxP-fabp7a-cerulean)* transgenic line was generated using the Tol2/Gateway system according to a previously published approach ^28^. The zebrafish full length *fabp7a* cDNA was RT-PCR amplified using a forward primer 5’- ATCGGAATTCTGGCCACCATGGTCGATGCATTTTGTGCCACTTGGG-3’ and a reverse primer 5’ -ATCGGCGGCCGCTGCCTTCTCGTATGTGCGCACGGCCTG-3’ introduced with an EcoRI and a NotI cut sites. The resultant PCR fragment was digested with EcoRI and NotI and inserted into the pENTRI1-loxP-mCherry-stop-loxP vector to generate pENTRI-loxP-mCherry-stop-loxP-fabp7a, which was then recombined with p5E-βactin2, p3E-cerulean-polyA, and pDestTol2pA to obtain the final construct: Tg(βactin2: loxP-mCherry-stop-loxP-fabp7a-cerulean) using Gateway LR Clonase II Plus Enzyme (ThermoFisher). Founder zebrafish (F0) were identified based on positive mCherry fluorescence and genotyping PCR. F1 stable transgenic fish were obtained by outcrossing. The resultant *Tg(*β*actin2: loxP-mCherry-stop-loxP-fabp7a- cerulean)* fish were then further outcrossed with the *Tg(cmlc2:creERT2)* fish to generate double transgenic fish, which were then incubated with 1 µM4-HT in system water for 24h to induce loxP-Cre recombination.

### TFEB transcription factor binding prediction

The Fabp7a promoter sequence was obtained from the Ensembl genome browser and exported to a BED format file including variation features with a total of 130 open peaks. Two different versions of the TFEB frequency matrix (MA0692.1, MA0692.2) were obtained from Jaspar, an open-access database of eukaryotic transcription factor binding profiles (PMID 14681366). The detection threshold for each frequency matrix was calculated based on the log-odds probabilities for the nucleotide in each position: 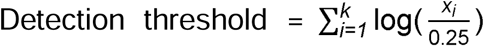 where *i, x,* and *k* represent nucleotide position, the probability of the major nucleotide in each position, and the last nucleotide position, respectively. TFEB enriched motifs in the Fabp7a promoter sequence were identified using HOMER with the danRer11 version of the zebrafish genome as a reference (PMID 20513432).

### Statistics

The unpaired two-tailed student’s *t*-test was used to compare two groups. One-way analysis of variance (ANOVA) was used to assess differences among three or more groups. For the animal survival rate comparison, the log-rank test was used to determine the difference. For all dot plots, each value represents the mean ± standard error (SE). All statistical analyses were performed with GraphPad Prism 7. For the post hoc analysis, Tukey’s test was employed to confirm the findings.

## Author contributions

Conceptualization: Y.D., X.L., and X.X.; Methodology: Y.D., F.Y.; Software: Y.D., F.Y. and Y.Z.; Validation: Y.D. and X.X.; Formal analysis: Y.D., F.Y., Y.Z., D.M.R. and X.X.; Investigation: Y.D. and X.X.; Data curation: Y.D., F.Y., B.Y., W.W., and D.M.R. Writing - original draft: Y.D. and X.X.; Writing - review & editing: Y.D., B.Y., and X.X.; Supervision: X.L. and X.X.; Project administration: X.X.; and Funding acquisition: X.X.

## Acknowledgments

This study was supported in part by grants from the NIH (HL107304 and HL081753) and the Mayo Foundation to X.X. We thank Beninio Gores and Briana M. Skufca for managing the zebrafish facility.

