## Supplemental Materials for "An mTOR-Tfeb-Fabp7a signaling axis can be harnessed to ameliorate *bag3* cardiomyopathy in adult zebrafish"

**Supplemental Figure 1. The *bag3* cardiomyopathy model manifested accelerated cardiac aging phenotypes**

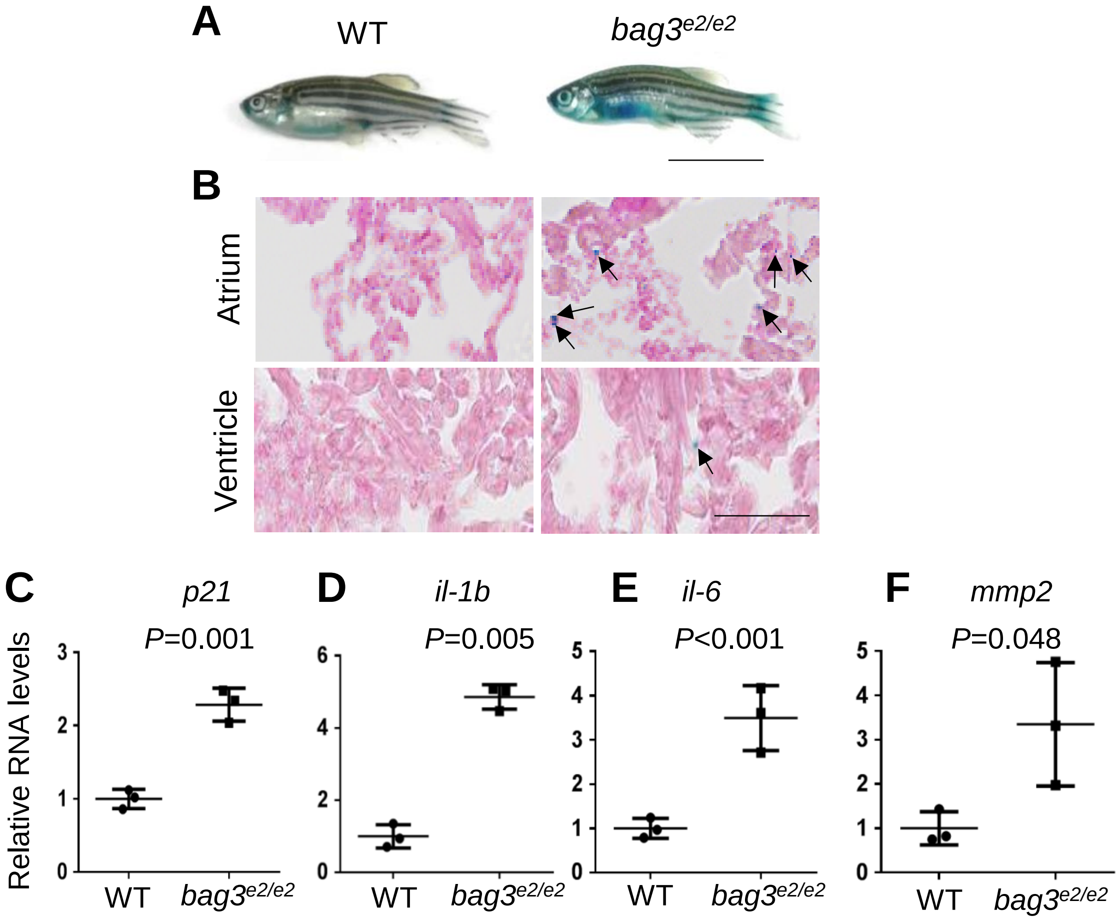


**A-B**, Representative images of senescence-associated β-galactosidase activity (SA-β-gal) staining in the whole fish body (A) and in the heart chamber (B) at 6 months of age. Arrows point to positive SA-β-gal staining signals. **C-D**, Quantitative RT-PCR analysis of senescence marker p21 and SASPs. n=3 biological replicates, one-way ANOVA. Scale bars in A, 1 cm, in B, 100 μm.

### Supplemental Figure 2. Volcano plots representation of differential expression and IPA pathway analysis

**A**

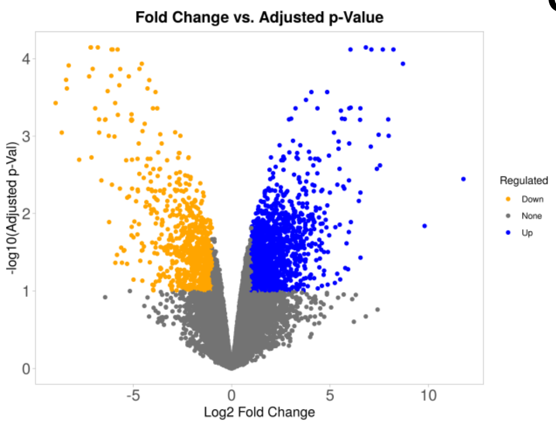


**C**

| Ingenuity Canonical Pathways | -log(p-value) |
|---|---|
| Cell Cycle Checkpoints | 25.9 |
| Mitotic Prometaphase | 20.3 |
| Activation of the pre-replicative complex | 17.8 |
| Kinetochore Metaphase Signaling Pathway | 16.4 |
| Synthesis of DNA | 16.3 |
| Cell Cycle Control of Chromosomal Replication | 13.6 |
| RHO GTPases Activate Formins | 13.4 |
| Mitotic Metaphase and Anaphase | 12.5 |
| Mitotic G1 phase and G1/S transition | 11.6 |
| Hereditary Breast Cancer Signaling | 9.53 |

**B**

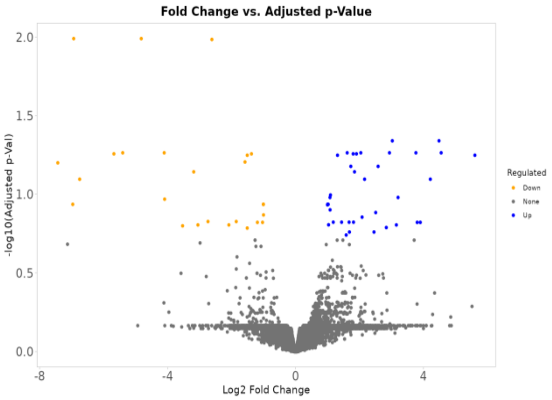


**D**

| Ingenuity Canonical Pathways | -log(p-value) |
|---|---|
| Circadian Clock | 3.61 |
| Protein folding | 2.96 |
| Macrophage Alternative Activation Signaling Pathway | 2.15 |
| NR1D1 (REV-ERBA) represses gene expression | 2.09 |
| NGF processing | 2.09 |
| PRPP Biosynthesis I | 2.09 |
| Oxidized GTP and dGTP Detoxification | 2.09 |
| Transmission across Electrical Synapses | 1.99 |
| Glycogen Biosynthesis II (from UDP-D-Glucose) | 1.84 |
| Unfolded protein response | 1.83 |

**A-B**, Volcano plots representation of differential expression in the *mtor*$^{xu015/+}$ mutant (A) and the *Tg(cmlc2:tfeb)* transgenic hearts (B). **C-D,** The top 10 DE gene enriched pathay suggested by IPA in the *mtor*$^{xu015/+}$ mutant (C) and the *Tg(cmlc2:tfeb)* transgenic hearts (D).

**Supplemental Figure 3. The TFEB transcription factor is predicted to bind to the Fabp7a promoter**

**A**

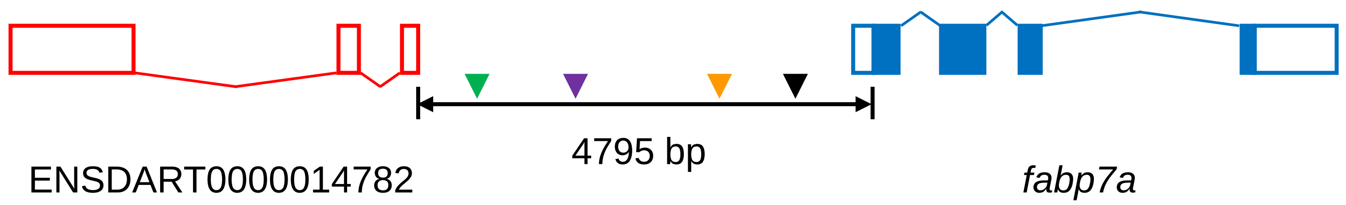


**B**

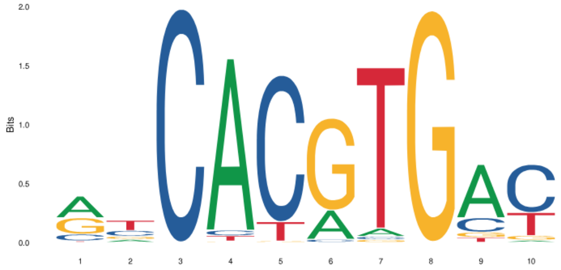


MA0692.1

| | A | C | T | G |
|---|---|---|---|---|
| **1** | 0.468099 | 0.146358 | 0.353218 | 0.032325 |
| **2** | 0.134464 | 0.260791 | 0.144291 | 0.460453 |
| **3** | 0.001946 | 0.99771 | 0 | 0.000343 |
| **4** | 0.933383 | 0.030631 | 0.00664 | 0.029346 |
| **5** | 0 | 0.875615 | 0.008339 | 0.116045 |
| **6** | 0.234664 | 0.024469 | 0.740442 | 0.000425 |
| **7** | 0.04785 | 0.017812 | 0.015809 | 0.918529 |
| **8** | 0.001029 | 0.001144 | 0.996569 | 0.001258 |
| **9** | 0.684281 | 0.183888 | 0.068467 | 0.063364 |
| **10** | 0.029615 | 0.616262 | 0.058393 | 0.29573 |

Detection threshold = 4.721592

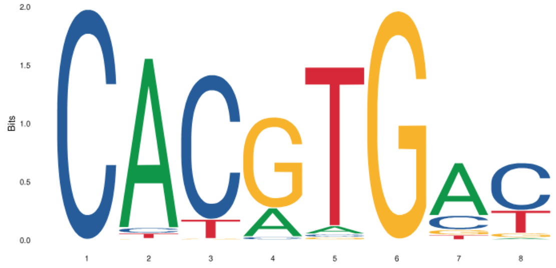


MA0692.2

| | A | C | G | T |
|---|---|---|---|---|
| **1** | 0.001946 | 0.99771 | 0 | 0.000343 |
| **2** | 0.933383 | 0.030631 | 0.00664 | 0.029346 |
| **3** | 0 | 0.875615 | 0.008339 | 0.116045 |
| **4** | 0.234664 | 0.024469 | 0.740442 | 0.000425 |
| **5** | 0.04785 | 0.017812 | 0.015809 | 0.918529 |
| **6** | 0.001029 | 0.001144 | 0.996569 | 0.001258 |
| **7** | 0.684281 | 0.183888 | 0.068467 | 0.063364 |
| **8** | 0.029615 | 0.616262 | 0.058393 | 0.29573 |

Detection threshold = 4.18395

**C**

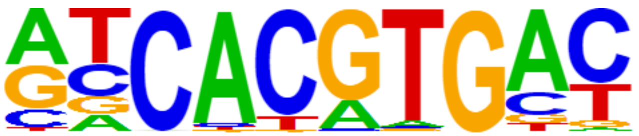

TFEB (MA0692.1)
p-value = $1e^{-4}$

**A,** Schematic of *fabp7a* promoter sequence. Colorless and colored rectangles represent untranslated and translated exon, respectively. Inverted triangles represent potential TFEB binding sites, with each color of the triangle representing a different sequence: GTCACGTGAT (green), GACATGTGGT (purple), GGCATGTGAC (orange), and GTCACGTGTA (black). **B,** Two different versions of TFEB binding motifs and their corresponding frequency matrices with detection thresholds **C,** Enriched TFEB binding motifs in *fabp7a* promoter sequence

**Supplemental Table 1.** List of 15 overlapping genes downregulated in both the *mtor*$^{xu015/+}$ haploinsufficiency mutant and the *Tg(cmlc2-tfeb)* transgenic hearts

| Gene | Locus | *mtor*$^{xu015/+}$ vs WT log2(FC) | *Tg(cmlc2-tfeb)* vs WT log2(FC) |
|---|---|---|---|
| *bscl2l* | chr14:6165398-6181950 | -1.1 | -2.4 |
| *smtnl1* | chr14:14938412-14954153 | -1.3 | -1.5 |
| *fabp7a* | chr17:15274756-15277126 | -1.6 | -2.3 |
| *ankrd9* | chr17:29335653-29336841 | -1.4 | -1.4 |
| *si:dkey-238c7.13* | chr18:7047760-7050665 | -1.1 | -2.3 |
| *kcnj5* | chr18:47136468-47207497 | -2.2 | -1.2 |
| *edn1* | chr19:3938958-3944172 | -1.1 | -1.0 |
| *ENSDARG00000020455* | chr19:11036022-11048001 | -1.3 | -4.9 |
| *grn1* | chr19:41146726-41165805 | -1.6 | -2.2 |
| *Novel gene* | chr19:18031738-18032158 | -1.5 | -4.2 |
| *MYADM (1 of 2)* | chr2:37470848-37480076 | -1.1 | -1.3 |
| *mpz* | chr2:44320982-44344524 | -1.7 | -1.7 |
| *CIDEC* | chr6:22886995-22896421 | -1.0 | -2.7 |
| *Novel gene* | chr7:53207421-53212375 | -1.2 | -6.3 |
| *CABZ01079442.1* | chr9:5289139-5330274 | -1.3 | -3.6 |
